## Supplementary figures and images for "A community-science approach identifies genetic variants associated with three color morphs in ball pythons (*Python regius*)"

### S1 Fig

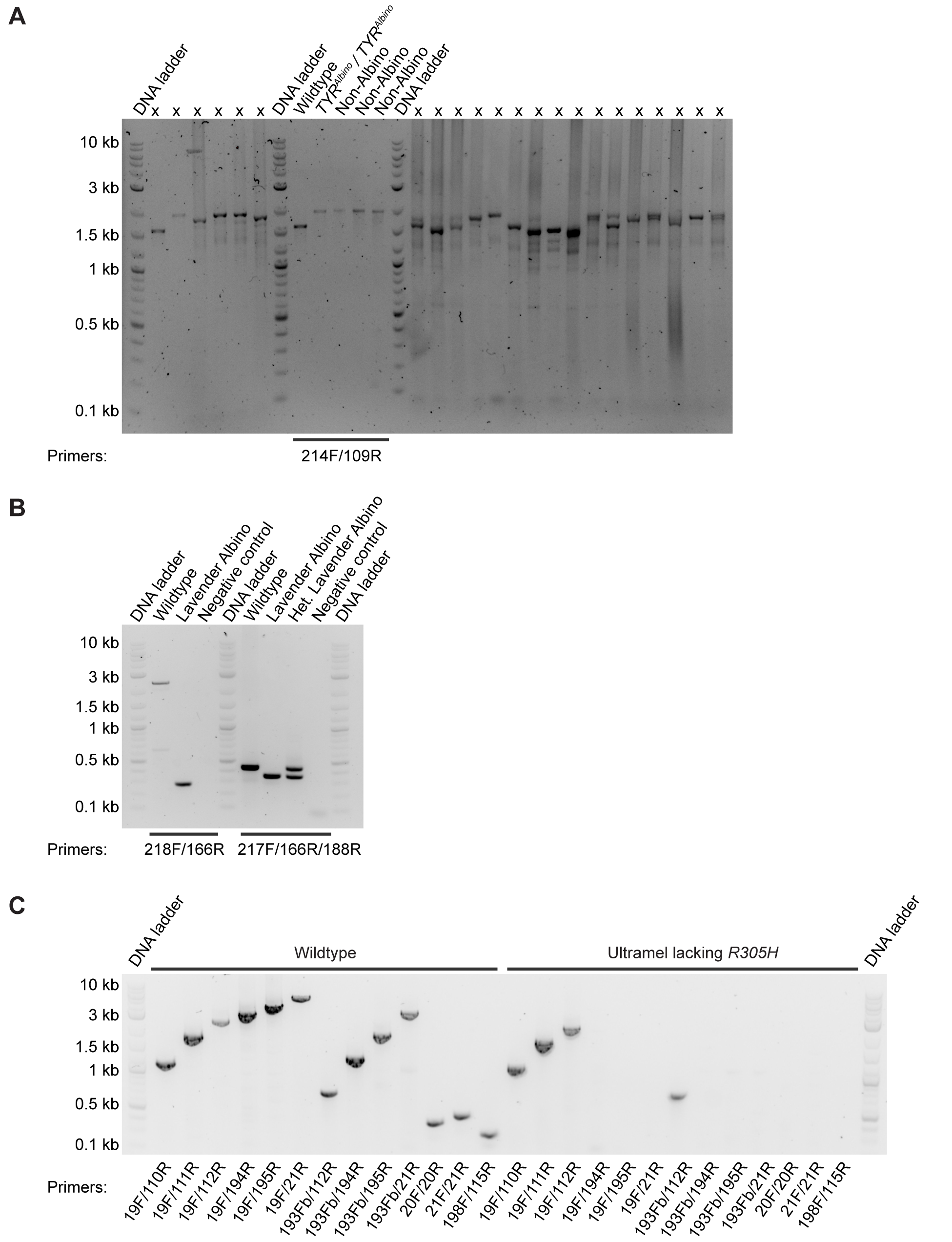
