## Supplementary material for "A community-science approach identifies genetic variants associated with three color morphs in ball pythons (*Python regius*)": S1 Table

**S1 Table. Variants used in the Albino association study and primers to genotype these variants.**

| **Gene** | **Variants and PCR primers** | **Annealing temperature** | **Sequencing primer** |
| --- | --- | --- | --- |
| *TYR* | Variants downstream of coding region 1:  5’-CTT CAT GGC AAG TAT [G/C]-3’  5’-TTG TAT GGT ACA TGC [A/C]-3’  Primers:  101F 5'-GACATCCCAGCAAA CAAAC-3' 101R 5'-ATACACGCATCACAGGACCA-3' | 61°C | 101F |
|  | Variant upstream of coding region 4:  5’-AAT ATT AAT ATC GAA [A/C]-3’  Variants downstream of coding region 4:  5’-AAA CAT CTA AAA CAG [A/T]-3’  5’-CAT ACT CAT CAT GCA [A/G]-3’  5’-TTG ATT TTG AGG GGG [C/T]-3’  Primers:  14F 5'-AGCTGATCTCCTGCATCTGG-3' -3' 94R 5'-AAAAGTGCCAAGGAAGCTGA-3' | 57°C | 14F |
| *TYRP1* | Variants downstream of coding region 6:  5’-ACT TCC AGA ATA TAA [CAA/---]-3’  5’-GCT TAT GAC TTT CGA [C/T]-3’  Primers:  96F 5'-CCAGAAAACCTGGGCTACAG-3' 96R 5'-ACCAGGATCAAAGCAGCAAC-3' | 61°C | 96F |
| *TYRP2* | Variants upstream of coding region 3:  5’-AAT CTA TAG CCT CTG [A/G]-3’  5’-ATG GGG GGG TTA AAT [G/-]-3’  Primers:  97F 5'-TGGCTTCATTATTATTCAGTCAGA-3' 102R 5'-CCTGAAAGCAGATGAGAGGAA-3' | 57°C | 102R |
| *OCA2* | Variant downstream of coding region 21:  5’-GTC CTT TGT CTT GGC [A/G]-3’  Primers:  47F 5'-GCAGAATTTTAATTTGAGTTATAGCA-3' 47R 5'-GCAATAAAGTAACTGGAGATGCAGA-3' | 57°C | 47F |
|  | Variant within coding region 24:  5’-CCT TCT TGT TGC TCA [C/T]-3’  Primers:  87F 5'-GAAAGTGCTGAGTCCTTGCTT-3' 87R 5'-TCAAAAATTTAACAAAAGCTGGAA-3' | 52°C | 87F |
