## Supplementary material for "A community-science approach identifies genetic variants associated with three color morphs in ball pythons (*Python regius*)": S2 Table

**S2 Table. Conservation of gene structure of melanogenesis genes.**

| **Gene** | **Number of coding regions** | **Protein sequences used in multi-species alignment** | **Conservation of coding-region boundaries relative to protein sequences** |
| --- | --- | --- | --- |
| *TYR* | 5 | Mouse (*NP_035791.1*) Chicken (*NP_989491.1*) Anole lizard (*XP_003219419.1*) | Perfectly conserved across species |
| *TYRP1* | 7 | Chicken (*NP_990376.2*) Anole lizard (*XP_008101333.1*) Corn snake (*XP_034266320.1*) | Perfectly conserved across species |
| *TYRP2* | 8 | Chicken (*NP_990266.1*) Anole lizard (*XP_003218710.1*) Corn snake (*XP_034273310.1*) | Perfectly conserved across species |
| *OCA2* | 24 | Chicken (*XP_025002402.1*) Anole lizard (*XP_008105312.1*) Corn snake (*XP_034287267.1*) | Perfectly conserved across species |
| *SLC7A11* | 12 | Chicken (*XP_426289.3*) Anole lizard (*XP_003221726.1*) Corn snake (*XP_034257397.1*) | Perfectly conserved across species |
| *SLC24A5* | 9 | Mouse (*NP_778199.2*) Chicken (*NP_001033586.2*) Anole lizard (*XP_008119677.1*) | Perfectly conserved across species |
| *SLC45A2* | 7 | Mouse (*NP_444307.1*) Chicken (*NP_001076833.2*) Corn snake (*XP_034298386.1*) | Conserved across species, except for the boundary between coding regions 6 and 7, which is shifted by two amino acids in mouse relative to chicken and corn snake |
