## Supplementary material for "A community-science approach identifies genetic variants associated with three color morphs in ball pythons (*Python regius*)": S3 Table

**S3 Table. Primers to amplify and sequence coding regions of melanogenesis genes.**

| **Gene** | **Coding region(s)** | **Primers** | **Annealing temperature** | **Sequencing primer(s)** |
| --- | --- | --- | --- | --- |
| *TYR* | 1 | 12F 5'-AGCCTGTGAGACGGACAAAA-3' 12R 5'-TCAGAAACCATCTCCACATCC-3' | 57°C | 12F and 12R |
|  | 2 | 88F 5'-TCTCCTTATCTGGGCTGGAA-3' 88R 5'-ATGAGGGGTAATTGCTCCAA-3' | 52°C | 88F |
|  | 3 | 13F 5'-ACTTTCAGGTGGGCAGCAG-3' 13R 5'-GCTGACAACTAAAATCTCTGCAA-3' | 52°C | 13F |
|  | 4 | 14F 5'-AGCTGATCTCCTGCATCTGG-3' 14R 5'-TCGGTGGGCTCTTATTTCTG-3' | 57°C | 14F |
|  | 5 | 15F 5'-CTGATAGGCTCCCATCCAAG-3' 15R 5'-GGAAAATATTGTACAAACAGGAACA-3' | 57°C | 15F |
| *TYRP1* | 1 | 16F 5'-CAGGAGTTTTACATCCTTCTTATTTC-3' 16R 5'-TCCATTCTTCAAACATCAGCA-3' | 52°C | 16F |
|  | 2 and 3 | 17F 5'-TGATGAAGCATGATTTGAATTTG-3' 17R 5'-TTTAAGCCGGCTAGATTCCA-3' | 57°C | 17F and 17R |
|  | 4 | 18F 5'-GCTCTTTTCTCTAAGTCTGACCTC-3' 18R 5'-TCTTGTCCCACAAAAGGATTT-3' | 57°C | 18F |
|  | 5 | 19F 5'-TTTTGTGCAATGTTATGTTAGGC-3' 19R 5'-TGGGATCTTGGCATCTTCTT-3' | 57°C | 19F |
|  | 6 | 20F 5'-GCATTGTTTTATCAGCCATGAA-3' 20R 5'-GCAGCTGCAAAAAGGAGTACA-3' | 57°C | 20F |
|  | 7 | 21F 5'-CATACTGAAATTCTTGGGACTCA-3' 21R 5'-GGAATTGAGACAAATCCTTGG-3' | 57°C | 21F |
| *TYRP2* | 1 | 22F 5'-GTCAGTTCGAATGGGGCTTG-3' 22R 5'-GAACAGTGACCAGCCTAGCA-3' | 57°C | 22F |
|  | 2 | 23F 5'-CAGGGTAGCGTCTAGCTTTG-3' 23R 5'-TGATCTAAAGGGCCATTGGT-3' | 57°C | 23F |
|  | 3 | 24F 5'-TGGTAGAGAGGACCCAGAGG-3' 24R 5'-TCCTGTAATGAGAATCCCTTCC-3' | 57°C | 24F |
|  | 4 | 25F 5'-GAAACATTGCAAAGATATGAAGGA-3' 25R 5'-CATCTGTATTGATTAGGCAGTTGAA-3' | 57°C | 25F |
|  | 5 and 6 | 26F 5'-TCAATGTGCTATTAATTATTCAGGTAA-3' 26R 5'-AAAGGTATTATTTCCATTTTGTGC-3' | 57°C | 26F |
|  | 7 | 27F 5'-AGACAATGGCACATGGTGAA-3' 27R 5'-AGCTGCAGGCTAATCCCTTG-3' | 57°C | 27F |
|  | 8 | 28F 5'-GCTGTGGAAGGGTTCAGAAG-3' 28R 5'-GGAGTTGAAGTCCACGCATC-3' | 57°C | 28R |
| *OCA2* | 1 and 2 | 82F 5'-CTTTAACTGAACATACCAATGGAG-3' 82R 5'-TGTGGATTGCTGAGTCAAAGA-3' | 57°C | 82F |
|  | 3 | 30F 5'-TTGCTTGTCAGTTCTAATATAAATCC-3' 30R 5'-TCTATCTAGCATTTCCTTCTCCAA-3' | 57°C | 30F |
|  | 4 | 31F 5'-TGTCACATTCACTAACAGATACATTCT-3' 31R 5'-TCTCTATTGCACTGGGATGTTC-3' | 52°C | 31F |
|  | 5 | 83F 5'-TTGTGTTATGAGTGGGTGAAGC-3' 32R 5'-GCCGGACCTAGGAATCATAAA-3' | 57°C | 32R |
|  | 6 | 33F 5'-CATATTTTTCCGGAATCACGA-3' 33R 5'-GCAAACAAAATATAAGCCATTTCA-3' | 57°C | 33F |
|  | 7 | 34F 5'-CAGTATAATGAGAAAGGGGGATG-3' 34R 5'-TGTGCACACTTTCCTTGGAG-3' | 57°C | 34F |
|  | 8 | 35F 5'-TGGAGTCTTTTCTATCAAATTAGCAA-3' 35R 5'-TGGCAGCACATAAAGTGGTC-3' | 57°C | 35F |
|  | 9 | 36F 5'-CAGAGTAATAGCAGCAAGATAAATAGC-3' 36R 5'-ACTTAACAGAAGTGTGCTGATTCTT-3' | 57°C | 36F |
|  | 10 | 37F 5'-TCTTTGTATCCCACTGAAGACC-3' 37R 5'-CAAAGACCCCTGCTGTTCAT-3' | 57°C | 37F |
|  | 11 | 161F 5'-TTTGAATTTCCAAGACTCAATGAC-3' 161R 5'-GCTGAAGCAGTTTAAAAGGGAAC-3' | 57°C | 161R |
|  | 12 | 38F 5'-GCATGCTCAATTTGATCTGAA-3' 38R 5'-GCAGAACTGAAGCAATTAACCTG-3' | 57°C | 38F |
|  | 13 | 85F 5'-TGTTGATCACCTTCACCTATTTCA-3' 85R 5'-ACGCTATTTTGATTCCATTCAT-3' | 57°C | 85R |
|  | 14 | 40F 5'-AGGCTCTGCTTTGTTGTTAAGT-3' 40R 5'-AGTGCCAGAATGGCTGTGTT-3' | 57°C | 40F |
|  | 15 | 41F 5'-AGAGACATGGTGTTAATTCTGCT-3' 41R 5'-GAAGCTGACTATAAATCCAATCCA-3' | 57°C | 41F |
|  | 16 | 42F 5'-GCCTGATTTGGTTGTAGTATTGC-3' 42R 5'-TTGCTGAGGCTGAGAAACAC-3' | 57°C | 42F |
|  | 17 | 43F 5'-TGTGTCATGACTAATGGTAAGTGG-3' 43R 5'-TGTGTCCTTGGCAGATTTCA-3' | 57°C | 43F |
|  | 18 | 44F 5'-TCACATTTGCTTCTGACTTGG-3' 44R 5'-GTCCCTGAAGCAGTGGTGTT-3' | 57°C | 44F |
|  | 19 | 45F 5'-AACTATGATTAGGTGCTACTCCACA-3' 45R 5'-AAGAGCTCCTAACTCACTCACACTC-3' | 57°C | 45F |
|  | 20 | 46F 5'-GCAAGTGGGCAATTTTGTCT-3' 46R 5'-GGCTTGCCTTTGAATCTAGG-3' | 57°C | 46R |
|  | 21 | 47F 5'-GCAGAATTTTAATTTGAGTTATAGCA-3' 47R 5'-GCAATAAAGTAACTGGAGATGCAGA-3' | 57°C | 47F |
|  | 22 | 48F 5'-CACTCATTTCGTCCCAAGGT-3' 48R 5'-TGACTGAAAAATTAGAAATCACAACC-3' | 57°C | 48F |
|  | 23 | 49F 5'-GATGACTCTCCTTTCCCCATT-3' 49R 5'-TTTTGCCATTGAAATGGAGTC-3' | 57°C | 49F |
|  | 24 | 87F 5'-GAAAGTGCTGAGTCCTTGCTT-3' 87R 5'-TCAAAAATTTAACAAAAGCTGGAA-3' | 52°C | 87R |
| *SLC7A11* | 1 | 71F 5'-TAGGCAGGAAGGGAGTGTGT-3' 71R 5'-CCAACCAAACTTTTGATCTTCA-3' | 57°C | 71F |
|  | 2 | 143F 5'-TTGGCAGAAGATTAGTTAGGACAA-3' 141R 5'-TGACAGCATCAAAATAAGTTGTGT-3' | 57°C | 143F |
|  | 3 | 109F 5'-GCTGCAATTTTACTTGTGATGG-3' 109R 5'-TCCAAATCAAACCAATTTTTCA-3' | 57°C | 109F |
|  | 4 | 74F 5'-TTTTGCTAATGCCTGGAATG-3' 74R 5'-AAATTGAACAAGCAGAGGCTAA-3' | 57°C | 74F |
|  | 5 | 75F 5'-TGGGGCAGATAAATAAACTGAA-3' 75R 5'-TTCCCATCCCAAACTTAGCA-3' | 52°C | 75F |
|  | 6 | 76F 5'-GCCTCACCTCCTATACCACA-3' 76R 5'-ACAACCTGACCATCATGCAA-3' | 57°C | 76F |
|  | 7 and 8 | 77F 5'-CAATCAGATGCCATGTAACAAGA-3' 77R 5'-TGTCAGTGATATTTACATTTGGAGAA-3' | 57°C | 77F |
|  | 9 | 78F 5'-CTTCTGGTGCTTGCCTTACA-3' 78R 5'-CCCCTCCAAACATAAAATGG-3' | 57°C | 78F |
|  | 10 | 79F 5'-AAACTTGAACTTGGCTGTTATGG-3' 79R 5'-TGAGCAACCAGAAAAGAGGA-3' | 57°C | 79F |
|  | 11 | 80F 5'-GCACTGGTCTCCCTGAATATC-3' 80R 5'-AAGATGCCTTACTATGACAGTGAGAA-3' | 52°C | 80F |
|  | 12 | 81F 5'-GGTCATAGTTGTATTCACATTGATCT-3' 81R 5'-GCCCCAATTTAACTTCTCCA-3' | 57°C | 81F |
| *SLC24A5* | 1 | 220F 5'-GACGAAGAGCGCTGAAGAAC-3' 226R 5'-TTCCCACCCATTAGCAGAAC-3' | 57°C | 220F and 226R |
|  | 2 | 221F 5'-GGAGATGGGTGGCATACAAA-3' 58R 5'-CTAAACTGCAGTCGCATTGTG-3' | 60°C | 58R |
|  | 3 and 4 | 59F 5'-TGCTTGGAGTCACTAGAAATGG-3' 59R 5'-TTTTTGGTACCCCCAGTTGA-3' | 52°C | 59F |
|  | 5 | 60F 5'-GGAGACTGCTGCACTAGATGG-3' 60R 5'-TGCTTTCTGTCCAGTTTTATCTGA-3' | 57°C | 60F |
|  | 6 | 105F 5'-CGCCAATGGAAAACATCATT-3' 105R 5'-TGCCTTGTAATTTGGGGTTT-3' sequencing only: 61R 5'-GCAGCTCTTGCAACACTTTCT-3' | 57°C | 105F, 105R, and 61R |
|  | 7 | 62F 5'-TGCACATCAGGATCTCATTTCT-3' 62R 5'-TCGGGCATTAACTCTTAACCTC-3' | 57°C | 62R |
|  | 8 and 9 | 106F 5'-TGTGACCCCCTGAAATATCACT-3' 106R 5'GAGACTGACCCTTTCATTCCTT--3' | 57°C | 106F and 106R |
| *SLC45A2* | 1 | 64F 5'-AACAGCCTCCTCGATACCAG-3' 64R 5'-CCTTGCCAACTAATCTATTGCAG-3' | 57°C | 64F |
|  | 2 | 65F 5'-TGCCAGTTGAATTACATGCAC-3' 65R 5'-TGCTTGCAACTGATGCTCTG-3' | 57°C | 65F |
|  | 3 | 66F 5'-AACATTTGCCAAACGGACTT-3' 66R 5'-TTTGGACCACGTGACATGAA-3' | 52°C | 66F |
|  | 4 | 107F 5'-GGGCGGCATAAACATTTAAC-3' 107R 5'-CACAGGGTTGGCTCAGAGG-3' | 57°C | 107F |
|  | 5 | 68F 5'-TTTGCATTCCCAATTGTCTG-3' 68R 5'-TCCCTGAGTTAATGAGTGTGAAA-3' | 52°C | 68R |
|  | 6 | 69F 5'-TGAGAAACACTGAACTGAAACATC-3' 69R 5'-TGGACTGGAGTCCTTCCTTT-3' | 57°C | 69F |
|  | 7 | 70F 5'-CGGTGTGCTTTAGCTGTGTG-3' 70R 5'-CAATGGAAGAAAAAGAATTGTCA-3' | 52°C | 70F |
