## Supplementary material for "A community-science approach identifies genetic variants associated with three color morphs in ball pythons (*Python regius*)": S4 Table

**S4 Table. Primers to amplify non-coding regions of *TYR*.**

| **Gene** | **Region** | **Primers** | **Annealing temperature** | **Sequencing primer(s)** |
| --- | --- | --- | --- | --- |
| *TYR* | Promoter | 208F 5'-TTCTCGATGAGATGGAAGCA-3' 204R 5'-GCTGCCATTTGTCTTCCAAT-3' | 57°C | 208F and 204R |
|  |  | 139F 5'-ATGCCAGAAGATAGGCTCCA-3' 7R 5'-TTGCCATGAAGAGAAGAAGGA-3' | 57°C | 139F and 7R |
|  | Intron 1 | 101F 5'-GACATCCCAGCAAACCAAAC-3' 116R 5'-CCCCCAAGTAAAATTGAATAGA-3' | 57°C | 116R |
|  |  | 117F 5'-TCCCAAACTGCAGTGAAATG-3' 11R 5'-CCGGAAGCTGAAGTTAGCC-3' | 57°C | 117F |
|  | Intron 2 | 8F 5'-GGTTCTGGTCCCAACTCTGT-3' 154R 5'-AGGAGTCAAGACTCACTTGAAGG-3' | 57°C | 154R |
|  |  | 120F 5'-GGCCAAATGAGTTAATTTGCAT-3' 120R 5'-GCTGTCCCAGGATCATTGAG-3' | 57°C | 120R |
|  |  | 155F 5'-GTTTGGGGAAAGGCAATGTT-3' 155R 5'-CTGACTCCGAGGCACCAG-3' | 57°C | 155F |
|  |  | 121F 5'-TGCACTTAGGGTTATGCCATC-3' 123R 5'-GCAGAGTAGCCGTTGAATCC-3' | 57°C | 123R |
|  |  | 156F 5'-CCAGAAGAAACAGACACAACACC-3' 156 5'-TGGCCCTGATGACTAAAGAGA-3' | 57°C | 156F |
|  |  | 124F 5'-TGCCAATACATCTGCTGGTT-3' 125R 5'-GGTGAGGAAGTTGGAACAGG-3' | 57°C | 124F |
|  |  | 125F 5'-CTAGGCAAAAAGGGCTGAAA-3' 90R 5'-ATTGCAGTACTTGGGCTTGC-3' | 57°C | 90R |
|  | Intron 3 | 91F 5'-AACGATCCGGTCTTCATCC-3' 91R 5'-TAGGGCAATGACAGAGCACA-3' | 57°C | 91F |
|  |  | 209F 5'-AACCCATGGTGCACTTTACT-3' 209R 5'-GTGCCCACCCTGATGTTATT-3' | 60°C | 209F |
|  |  | 92F 5'-ACCACCCCAGTCACCCTATT-3' 157R 5'-TGGAACACTATTTGCACTGTTTC-3' | 57°C | 92F |
|  |  | 130F 5'-GGTCTTCTGTGAAGGCTGCT-3' 131R 5'-CCTCTTGACCTGCCTTTGTC-3' | 57°C | 130F |
|  |  | 158F 5'-TCCAAATCATACCCCAGAAA-3' 92R 5'-GCTAGAAACAAAGGGGCTGA-3' | 57°C | 158F |
|  |  | 134F 5'-GCGAGAGAGAGAGTTTTGCAT-3' 134R 5'-TTTCATCTTTGCCCTGTTCC-3' | 57°C | 134R |
|  |  | 159F 5'-AGGGGATTTCTGTCCTGTGA-3' 159R 5'-TGCTACGAATTAAGGGCACA-3' | 57°C | 149F |
|  |  | 135F 5'-CATTCCTGCAAAGAAACATCC-3'  14R 5'-TCGGTGGGCTCTTATTTCTG-3' | 57°C | 135F |
|  | Intron 4 | 14F 5'-AGCTGATCTCCTGCATCTGG-3' 137R 5'-TTCCTGAAACAGCATTGCAC-3' | 57°C | 137R |
|  |  | 137F 5'-ACCTCTACCCCTCCTGCACT-3' 138R 5'-CTGGAAAAACATGCAGCAAA-3' | 57°C | 137F |
|  |  | 138F 5'-CTGAGGGTCACAAGGAAAGG-3' 15R 5'-GGAAAATATTGTACAAACAGGAACA-3' | 57°C | 138F |
