## Supplementary material for "A community-science approach identifies genetic variants associated with three color morphs in ball pythons (*Python regius*)": S5 Table

**S5 Table. Primers to investigate the putative deletion in *TYRP1*.**

| **Gene** | **Region** | **Primers, polymerase, and annealing temperature**  (OneTaq, 57°C, unless specified) | **Amplification in Ultramels lacking *R305H*** |
| --- | --- | --- | --- |
| *TYRP1* | Coding region 5 and intron 5 | 19F 5’-TTTTGTGCAATGTTATGTTAGGC-3’  110R 5’-TCATGCTTGGATCTTGGTCA-3’ | Successful |
|  |  | 19F 5’-TTTTGTGCAATGTTATGTTAGGC-3’ 111R 5-GCCCAGCTCCCAACATTAGT-3’ | Successful |
|  |  | 19F 5’-TTTTGTGCAATGTTATGTTAGGC-3’ 112R 5’-TGGTGCATTGGAATATCTGC-3’ | Successful |
|  |  | 19F 5’-TTTTGTGCAATGTTATGTTAGGC-3’ 194R 5’-CCCTTCTTCCATGCAGTGAT-3’ | Failed |
|  |  | 19F 5’-TTTTGTGCAATGTTATGTTAGGC-3’ 195R 5’-AAATGGCTTTTCAGAGGAACT-3’ | Failed |
|  | Coding regions 5 to 7 | 19F 5’-TTTTGTGCAATGTTATGTTAGGC-3’ 21R 5’-GGAATTGAGACAAATCCTTGG-3’  Q5, 58°C | Failed |
|  | Intron 5 | 193Fb 5’-TGCATGTCACTCATGCCTCT-3’ 112R 5’-TGGTGCATTGGAATATCTGC-3’ | Successful |
|  |  | 193Fb 5’-TGCATGTCACTCATGCCTCT-3’ 194R 5’-CCCTTCTTCCATGCAGTGAT-3’  OneTaq, 60°C | Failed |
|  |  | 193Fb 5’-TGCATGTCACTCATGCCTCT-3’ 195R 5’-AAATGGCTTTTCAGAGGAACT-3’  OneTaq, 60°C | Failed |
|  | Intron 5 to coding region 7 | 193Fb 5’-TGCATGTCACTCATGCCTCT-3’ 21R 5’-GGAATTGAGACAAATCCTTGG-3’  OneTaq, 60°C | Failed |
|  | Coding region 6 | 20F 5’-GCATTGTTTTATCAGCCATGAA-3’ 20R 5’-GCAGCTGCAAAAAGGAGTACA-3’ | Failed |
|  | Coding region 7 | 21F 5’-CATACTGAAATTCTTGGGACTCA-3’ 21R 5’-GGAATTGAGACAAATCCTTGG-3’ | Failed |
|  | Downstream of coding region 7 | 198F 5’-TGGAAGATGGTGCAGAAATTA-3’ 115R 5’-GGGAGGCAAGATGTTCCATA-3’ | Failed |
