## Supplementary material for "A community-science approach identifies genetic variants associated with three color morphs in ball pythons (*Python regius*)": S6 Table

**S6 Table. Genotypes.**

| **Morph** | **ID** | **Genotype**  nd, not determined | | | | | | | | | | | | | | | | | |
| --- | --- | --- | --- | --- | --- | --- | --- | --- | --- | --- | --- | --- | --- | --- | --- | --- | --- | --- | --- |
|  |  | **Association study variants** | | | | | | | | | | | | **Albino**  **variants** | | | **Lavender**  **Albino**  **variant** | **Ultramel**  **variants** | |
|  |  | *TYR* | | | | | | *TYRP1* | | *TYRP2* | | *OCA2* | | *TYR* | | | *OCA2* | *TYRP1* | |
|  |  | 101F | | 14F | | | | 96F | | 102R | | 47F | 87F |  |  |  |  |  |  |
| Non, Non-Albino and  Non-Lavender Albino and  Non-Ultramel |  | CTT CAT GGC AAG TAT [G/C] | TTG TAT GGT ACA TGC [A/C] | AAT ATT AAT ATC GAA [A/C] | AAA CAT CTA AAA CAG [A/T] | CAT ACT CAT CAT GCA [A/G] | TTG ATT TTG AGG GGG [C/T] | ACT TCC AGA ATA TAA [CAA/---] | GCT TAT GAC TTT CGA [C/T] | AAT CTA TAG CCT CTG [A/G] | ATG GGG GGG TTA AAT [G/-] | GTC CTT TGT CTT GGC [A/G] | CCT TCT TGT TGC TCA [C/T] | *D394G* | *P384L* | Haplotype *TYR^Albino^* | Deletion of coding region 18 | *R305H* | Coding regions 6 and 7 |
|  |  |  |  |  |  |  |  |  |  |  |  |  |  |  |  | Hom, homozygous  Het, heterozygous  None, non-carrier |  |  |  |
| Albino | 5 | C/C | A/A | A/A | A/A | G/G | C/C | CAA/CAA | T/T | A/A | G/G | A/A | C/C | +/+ | +/+ | Hom | +/+ | +/+ | Present |
| Albino | 32 | C/C | A/A | A/A | A/A | G/G | C/C | CAA/CAA | C/C | G/G | G/- | A/G | C/T | +/+ | +/+ | Hom | +/+ | +/+ | Present |
| Albino | 52 | C/C | A/A | A/A | A/A | G/G | C/C | CAA/CAA | C/C | G/G | -/- | A/G | C/T | +/+ | +/+ | Hom | +/+ | +/+ | Present |
| Albino | 116 | C/C | A/A | A/A | A/A | G/G | C/C | CAA/--- | T/T | G/G | -/- | A/A | C/C | +/+ | +/+ | Hom | +/+ | +/+ | Present |
| Albino | 118 | C/C | A/A | A/A | A/A | G/G | C/C | CAA/CAA | T/T | A/G | G/G | A/A | C/C | +/+ | +/+ | Hom | +/+ | +/+ | Present |
| Albino | 183 | C/C | A/A | A/A | A/A | G/G | C/C | CAA/CAA | C/T | nd | nd | A/A | C/C | +/+ | +/+ | Hom | +/+ | +/+ | Present |
| Albino | 241 | C/C | A/A | A/A | A/A | G/G | C/C | CAA/CAA | C/C | A/G | G/- | A/A | C/C | +/+ | +/+ | Hom | +/+ | +/+ | Present |
| Albino | 258 | C/C | A/A | A/A | A/A | G/G | C/C | CAA/CAA | C/C | G/G | -/- | A/A | C/C | +/+ | +/+ | Hom | +/+ | +/+ | Present |
| Albino | 260 | C/C | A/A | A/A | A/A | G/G | C/C | CAA/CAA | C/C | G/G | -/- | A/G | C/T | +/+ | +/+ | Hom | +/+ | +/+ | Present |
| Albino | 272 | C/C | A/A | A/A | A/A | G/G | C/C | CAA/CAA | T/T | G/G | -/- | A/G | C/T | +/+ | +/+ | Hom | +/+ | +/+ | Present |
| Albino | 284 | C/C | A/A | A/A | A/A | G/G | C/C | --- | T/T | G/G | -/- | A/A | C/C | +/+ | +/+ | Hom | +/+ | +/+ | Present |
| Albino | 291 | C/C | A/A | A/A | A/A | G/G | C/C | nd | nd | A/A | G/G | A/A | C/C | +/+ | +/+ | Hom | +/+ | +/+ | Present |
| Albino | 303 | C/C | A/A | A/A | A/A | G/G | C/C | CAA/CAA | C/T | G/G | G/G | A/A | C/C | +/+ | +/+ | Hom | +/+ | +/+ | Present |
| Albino | Amel | C/C | A/A | A/A | A/A | G/G | C/C | CAA/CAA | T/T | A/G | G/G | A/A | C/T | +/+ | +/+ | Hom | +/+ | +/+ | Present |
| Albino | 98 | G/G | A/A | A/A | T/T | A/A | T/T | CAA | C/T | G/G | -/- | A/G | C/T | *D394G /  D394G* | +/+ | None | +/+ | +/+ | Present |
| Albino | 134 | G/G | A/A | A/A | T/T | A/A | T/T | CAA | C/C | G/G | G/- | A/A | C/C | *D394G /  D394G* | +/+ | None | +/+ | +/+ | Present |
| Albino | 1 | C/G | A/A | A/A | A/T | A/G | C/T | --- | T/T | G/G | -/- | A/A | C/C | *D394G / +* | +/+ | Het | +/+ | +/+ | Present |
| Albino | 29 | C/G | A/A | A/A | A/T | A/G | C/T | CAA/CAA | C/T | G/G | -/- | A/G | C/T | *D394G / +* | +/+ | Het | +/+ | +/+ | Present |
| Albino | 87 | C/G | A/A | A/A | A/T | A/G | C/T | CAA/CAA | T/T | G/G | G/G | A/A | C/T | *D394G / +* | +/+ | Het | +/+ | +/+ | Present |
| Albino | 127 | C/G | A/A | A/A | A/T | A/G | C/T | CAA/CAA | T/T | G/G | G/- | A/G | C/T | *D394G / +* | +/+ | Het | +/+ | +/+ | Present |
| Albino | 257 | C/G | A/A | A/A | A/T | A/G | C/T | CAA/--- | C/T | nd | nd | A/A | C/C | *D394G / +* | +/+ | Het | +/+ | +/+ | Present |
| Albino | 256 | G/G | A/A | C/C | T/T | A/A | T/T | CAA | T/T | G/G | G/G | A/A | C/T | +/+ | *P384L /  P384L* | None | +/+ | +/+ | Present |
| Albino | 91 | C/G | A/A | A/C | A/T | A/G | C/T | CAA | C/T | A/G | G/G | A/G | C/T | +/+ | *P384L / +* | Het | +/+ | +/+ | Present |
| Albino | 92 | C/G | A/A | A/C | A/T | A/G | C/T | CAA | T/T | G/G | -/- | A/A | C/C | +/+ | *P384L / +* | Het | +/+ | +/+ | Present |
| Albino | 162 | C/G | A/A | A/C | A/T | A/G | C/T | CAA | C/T | G/G | G/- | A/A | C/C | +/+ | *P384L / +* | Het | +/+ | +/+ | Present |
| Albino | 297 | C/G | A/A | A/C | A/T | A/G | C/T | CAA | C/C | G/G | G/- | A/G | T/T | +/+ | *P384L / +* | Het | +/+ | +/+ | Present |
| Albino | 302 | C/G | A/A | A/C | A/T | A/G | C/T | CAA/--- | T/T | G/G | G/- | A/A | C/C | +/+ | *P384L / +* | Het | +/+ | +/+ | Present |
| Albino | 329 | C/C | A/A | A/A | A/A | G/G | C/C | nd | nd | nd | nd | nd | nd | +/+ | +/+ | Hom | nd | nd | nd |
| Albino | 443 | C/C | A/A | A/A | A/A | G/G | C/C | nd | nd | nd | nd | nd | nd | +/+ | +/+ | Hom | nd | nd | nd |
| Albino | 457 | C/C | A/A | A/A | A/A | G/G | C/C | nd | nd | nd | nd | nd | nd | +/+ | +/+ | Hom | nd | nd | nd |
| Albino | 468 | C/G | A/A | A/C | A/T | A/G | C/T | nd | nd | nd | nd | nd | nd | +/+ | *P384L / +* | Het | nd | nd | nd |
| Albino | 484 | C/C | A/A | A/A | A/A | G/G | C/C | nd | nd | nd | nd | nd | nd | +/+ | +/+ | Hom | nd | nd | nd |
| Albino | 485 | G/G | A/A | C/C | T/T | A/A | T/T | nd | nd | nd | nd | nd | nd | +/+ | *P384L /  P384L* | None | nd | nd | nd |
| Albino | 495 | G/G | A/A | C/C | T/T | A/A | T/T | nd | nd | nd | nd | nd | nd | +/+ | *P384L /  P384L* | None | nd | nd | nd |
| Albino | 499 | C/G | A/A | A/A | A/T | A/G | C/T | nd | nd | nd | nd | nd | nd | *D394G / +* | +/+ | Het | nd | nd | nd |
| Albino | 547 | C/C | A/A | A/A | A/A | G/G | C/C | nd | nd | nd | nd | nd | nd | +/+ | +/+ | Hom | nd | nd | nd |
| Albino | 556 | C/C | A/A | A/A | A/A | G/G | C/C | nd | nd | nd | nd | nd | nd | +/+ | +/+ | Hom | nd | nd | nd |
| Albino | 564 | C/C | A/A | A/A | A/A | G/G | C/C | nd | nd | nd | nd | nd | nd | +/+ | +/+ | Hom | nd | nd | nd |
| Albino | 604 | C/C | A/A | A/A | A/A | G/G | C/C | nd | nd | nd | nd | nd | nd | +/+ | +/+ | Hom | nd | nd | nd |
| Albino | 626 | C/G | A/A | A/A | A/T | A/G | C/T | nd | nd | nd | nd | nd | nd | *D394G / +* | +/+ | Het | nd | nd | nd |
| Albino | 665 | C/C | A/A | A/A | A/A | G/G | C/C | nd | nd | nd | nd | nd | nd | +/+ | +/+ | Hom | nd | nd | nd |
| Albino | 736 | C/C | A/A | A/A | A/A | G/G | C/C | nd | nd | nd | nd | nd | nd | +/+ | +/+ | Hom | nd | nd | nd |
| Albino | 870 | C/C | A/A | A/A | A/A | G/G | C/C | nd | nd | nd | nd | nd | nd | +/+ | +/+ | Hom | nd | nd | nd |
| Albino | 900 | C/C | A/A | A/A | A/A | G/G | C/C | nd | nd | nd | nd | nd | nd | +/+ | +/+ | Hom | nd | nd | nd |
| Albino | 1032 | G/G | A/A | A/C | T/T | A/A | T/T | nd | nd | nd | nd | nd | nd | *D394G / +* | *P384L / +* | None | nd | nd | nd |
| Albino | 1057 | G/G | A/A | C/C | T/T | A/A | T/T | nd | nd | nd | nd | nd | nd | +/+ | *P384L /  P384L* | None | nd | nd | nd |
| Albino | 1124 | C/G | A/A | A/C | A/T | A/G | C/T | nd | nd | nd | nd | nd | nd | +/+ | *P384L / +* | Het | nd | nd | nd |
| Albino | 1130 | C/C | A/A | A/A | A/A | G/G | C/C | nd | nd | nd | nd | nd | nd | +/+ | +/+ | Hom | nd | nd | nd |
| Albino | 1013 | C/C | A/A | A/A | A/A | G/G | C/C | nd | nd | nd | nd | nd | nd | +/+ | +/+ | Hom | nd | nd | nd |
| Albino | 1030 | C/G | A/A | A/A | A/T | A/G | C/T | nd | nd | nd | nd | nd | nd | *D394G / +* | +/+ | Het | nd | nd | nd |
| Lav. Albino | 296 | G/G | A/A | A/C | A/T | A/A | C/T | CAA/CAA | T/T | nd | nd | G/G | T/T | +/+ | +/+ | None | Deletion /  Deletion | +/+ | Present |
| Lav. Albino | 308 | G/G | A/A | C/C | A/T | A/G | T/T | CAA/CAA | C/C | G/G | -/- | G/G | T/T | +/+ | +/+ | None | Deletion /  Deletion | +/+ | Present |
| Lav. Albino | 314 | G/G | A/A | C/C | A/A | G/G | T/T | nd | nd | G/G | -/- | G/G | T/T | +/+ | +/+ | None | Deletion /  Deletion | +/+ | Present |
| Lav. Albino | 318 | G/G | A/C | A/A | A/T | A/G | T/T | CAA/CAA | T/T | G/G | -/- | G/G | T/T | +/+ | +/+ | None | Deletion /  Deletion | +/+ | Present |
| Lav. Albino | 327 | G/G | A/A | C/C | A/T | A/G | T/T | CAA/CAA | T/T | G/G | -/- | G/G | T/T | +/+ | +/+ | None | Deletion /  Deletion | +/+ | Present |
| Lav. Albino | 309 | nd | nd | nd | nd | nd | nd | nd | nd | nd | nd | nd | nd | nd | nd | nd | Deletion /  Deletion | nd | nd |
| Lav. Albino | 315 | nd | nd | nd | nd | nd | nd | nd | nd | nd | nd | nd | nd | nd | nd | nd | Deletion /  Deletion | nd | nd |
| Lav. Albino | 319 | nd | nd | nd | nd | nd | nd | nd | nd | nd | nd | nd | nd | nd | nd | nd | Deletion /  Deletion | nd | nd |
| Lav. Albino | 663 | nd | nd | nd | nd | nd | nd | nd | nd | nd | nd | nd | nd | nd | nd | nd | Deletion /  Deletion | nd | nd |
| Lav. Albino | 664 | nd | nd | nd | nd | nd | nd | nd | nd | nd | nd | nd | nd | nd | nd | nd | Deletion /  Deletion | nd | nd |
| Lav. Albino | 838 | nd | nd | nd | nd | nd | nd | nd | nd | nd | nd | nd | nd | nd | nd | nd | Deletion /  Deletion | nd | nd |
| Lav. Albino | 896 | nd | nd | nd | nd | nd | nd | nd | nd | nd | nd | nd | nd | nd | nd | nd | Deletion /  Deletion | nd | nd |
| Lav. Albino | 1283 | nd | nd | nd | nd | nd | nd | nd | nd | nd | nd | nd | nd | nd | nd | nd | Deletion /  Deletion | nd | nd |
| Lav. Albino | 1328 | nd | nd | nd | nd | nd | nd | nd | nd | nd | nd | nd | nd | nd | nd | nd | Deletion /  Deletion | nd | nd |
| Het.  Lav. Albino | 64 | nd | nd | nd | nd | nd | nd | nd | nd | nd | nd | nd | nd | nd | nd | nd | Deletion / + | nd | nd |
| Non | 8 | G/G | A/A | C/C | A/A | A/A | T/T | CAA/CAA | C/T | G/G | -/- | A/A | C/C | +/+ | +/+ | None | +/+ | +/+ | Present |
| Non | 9 | G/G | A/C | A/C | A/A | G/G | C/T | CAA/CAA | T/T | A/G | G/G | A/G | C/T | +/+ | +/+ | None | +/+ | +/+ | Present |
| Non | 12 | G/G | A/A | A/C | A/A | G/G | C/T | CAA/CAA | T/T | G/G | -/- | A/A | C/C | +/+ | +/+ | None | +/+ | +/+ | Present |
| Non | 22 | G/G | A/A | A/C | A/T | A/G | C/T | CAA/CAA | C/T | G/G | G/- | A/A | C/C | +/+ | +/+ | None | +/+ | +/+ | Present |
| Non | 26 | G/G | A/C | C/C | A/T | A/G | T/T | CAA/CAA | C/T | G/G | G/- | A/G | C/T | +/+ | +/+ | None | +/+ | +/+ | Present |
| Non | 33 | G/G | A/A | A/A | A/A | G/G | C/C | CAA/CAA | T/T | A/G | G/G | A/A | C/C | +/+ | +/+ | None | +/+ | +/+ | Present |
| Non | 41 | C/G | A/A | C/C | A/A | A/A | C/C | CAA/CAA | T/T | G/G | G/- | A/G | C/C | +/+ | +/+ | None | +/+ | +/+ | Present |
| Non | 45 | G/G | A/A | C/C | A/A | G/G | T/T | CAA/CAA | C/C | A/G | G/G | A/G | C/T | +/+ | +/+ | None | +/+ | +/+ | Present |
| Non | 49 | G/G | A/A | A/C | A/T | A/G | C/T | CAA/CAA | T/T | G/G | G/- | A/A | C/C | +/+ | +/+ | None | +/+ | +/+ | Present |
| Non | 53 | G/G | A/A | C/C | A/A | G/G | T/T | CAA/CAA | C/T | A/G | G/G | A/A | C/C | +/+ | +/+ | None | +/+ | +/+ | Present |
| Non | 54 | G/G | A/A | C/C | A/A | G/G | T/T | CAA/CAA | T/T | G/G | -/- | A/A | C/T | +/+ | +/+ | None | +/+ | +/+ | Present |
| Non | 55 | G/G | A/A | C/C | A/T | A/G | T/T | CAA/CAA | C/T | G/G | G/- | A/A | C/C | +/+ | +/+ | None | +/+ | +/+ | Present |
| Non | 58 | G/G | A/A | C/C | T/T | A/G | T/T | CAA/CAA | T/T | A/G | G/G | A/A | C/C | +/+ | +/+ | None | +/+ | +/+ | Present |
| Non | 60 | G/G | A/C | A/C | A/A | G/G | C/T | CAA/CAA | C/T | A/G | G/G | A/A | C/C | +/+ | +/+ | None | +/+ | +/+ | Present |
| Non | 65 | G/G | A/A | A/C | A/T | A/G | C/T | CAA/CAA | T/T | G/G | G/- | A/A | C/C | +/+ | +/+ | None | +/+ | +/+ | Present |
| Non | 67 | G/G | C/C | C/C | A/A | G/G | T/T | CAA/CAA | C/T | G/G | G/- | A/A | C/C | +/+ | +/+ | None | +/+ | +/+ | Present |
| Non | 71 | G/G | A/A | C/C | A/T | A/G | T/T | CAA/CAA | C/T | G/G | -/- | A/G | C/C | +/+ | +/+ | None | +/+ | +/+ | Present |
| Non | 72 | C/C | A/C | C/C | A/A | G/G | T/T | CAA/CAA | C/T | G/G | G/G | A/G | C/C | +/+ | +/+ | None | +/+ | +/+ | Present |
| Non | 94 | G/G | A/A | C/C | A/T | A/G | T/T | CAA/--- | T/T | G/G | G/- | A/G | C/C | +/+ | +/+ | None | +/+ | +/+ | Present |
| Non | 100 | G/G | A/A | A/C | A/A | G/G | C/T | CAA/CAA | C/C | A/G | G/G | G/G | C/T | +/+ | +/+ | None | +/+ | +/+ | Present |
| Non | 105 | G/G | A/A | C/C | A/T | A/G | T/T | CAA/CAA | T/T | A/G | G/G | A/A | C/C | +/+ | +/+ | None | +/+ | +/+ | Present |
| Non | 112 | G/G | A/A | A/C | A/A | G/G | C/T | CAA/CAA | C/T | G/G | -/- | A/A | C/C | +/+ | +/+ | None | +/+ | +/+ | Present |
| Non | 113 | G/G | A/C | C/C | A/A | G/G | T/T | CAA/CAA | C/T | G/G | G/G | A/G | C/T | +/+ | +/+ | None | +/+ | +/+ | Present |
| Non | 117 | G/G | A/C | C/C | A/T | A/G | T/T | CAA/CAA | C/C | G/G | -/- | A/G | C/C | +/+ | +/+ | None | +/+ | +/+ | Present |
| Non | 124 | G/G | A/C | C/C | A/T | A/G | T/T | CAA/CAA | T/T | G/G | G/G | A/G | C/C | +/+ | +/+ | None | +/+ | +/+ | Present |
| Non | 128 | G/G | A/A | A/A | A/A | G/G | C/C | CAA/CAA | C/T | A/G | G/G | A/G | C/C | +/+ | +/+ | None | +/+ | +/+ | Present |
| Non | 131 | G/G | A/A | A/C | A/T | A/G | C/T | CAA/CAA | T/T | G/G | G/G | A/A | C/C | +/+ | +/+ | None | +/+ | +/+ | Present |
| Non | 133 | G/G | A/C | C/C | A/T | A/G | T/T | CAA/CAA | C/C | A/G | G/G | A/G | C/T | +/+ | +/+ | None | +/+ | +/+ | Present |
| Non | 135 | G/G | A/A | A/C | A/A | G/G | C/T | CAA/CAA | C/T | G/G | -/- | G/G | T/T | +/+ | +/+ | None | +/+ | +/+ | Present |
| Non | 156 | G/G | A/A | A/C | A/T | A/G | C/T | CAA/--- | C/T | A/G | G/- | A/A | C/C | +/+ | +/+ | None | +/+ | +/+ | Present |
| Non | 167 | G/G | A/A | A/C | A/T | A/G | C/T | CAA/--- | T/T | G/G | G/- | A/G | C/T | +/+ | +/+ | None | +/+ | +/+ | Present |
| Non | 169 | G/G | A/A | C/C | A/T | A/G | T/T | CAA/CAA | C/T | G/G | G/- | A/G | C/C | +/+ | +/+ | None | +/+ | +/+ | Present |
| Non | 170 | G/G | A/C | C/C | A/A | G/G | T/T | CAA/CAA | T/T | G/G | G/G | A/A | C/C | +/+ | +/+ | None | +/+ | +/+ | Present |
| Non | 174 | G/G | A/A | A/C | A/T | A/G | C/T | CAA/CAA | T/T | G/G | -/- | A/A | C/C | +/+ | +/+ | None | +/+ | +/+ | Present |
| Non | 178 | G/G | A/A | A/C | A/T | A/G | C/T | CAA/CAA | C/T | G/G | -/- | A/A | C/C | +/+ | +/+ | None | +/+ | +/+ | Present |
| Non | 180 | G/G | A/C | A/C | A/A | G/G | C/T | CAA/CAA | C/C | A/G | G/- | A/A | C/C | +/+ | +/+ | None | +/+ | +/+ | Present |
| Non | 186 | G/G | A/A | C/C | A/T | A/G | T/T | CAA/CAA | C/C | G/G | G/- | A/G | C/T | +/+ | +/+ | None | +/+ | +/+ | Present |
| Non | 191 | G/G | A/A | A/A | A/A | G/G | C/C | CAA/CAA | C/T | A/G | G/- | A/G | C/T | +/+ | +/+ | None | +/+ | +/+ | Present |
| Non | 195 | G/G | A/C | A/A | A/A | G/G | C/C | CAA/CAA | C/C | A/G | G/- | A/G | C/T | +/+ | +/+ | None | +/+ | +/+ | Present |
| Non | 196 | G/G | A/A | C/C | A/T | A/G | T/T | CAA/CAA | C/C | G/G | G/G | A/G | C/T | +/+ | +/+ | None | +/+ | +/+ | Present |
| Non | 197 | G/G | A/A | C/C | A/T | A/G | T/T | CAA/CAA | T/T | G/G | -/- | A/A | C/C | +/+ | +/+ | None | +/+ | +/+ | Present |
| Non | 205 | G/G | A/A | A/C | A/T | A/G | C/T | CAA/CAA | T/T | A/A | G/G | A/G | C/T | +/+ | +/+ | None | +/+ | +/+ | Present |
| Non | 209 | G/G | A/A | C/C | A/T | A/G | T/T | nd | nd | G/G | G/- | A/A | C/C | +/+ | *P384L / +* | None | +/+ | +/+ | Present |
| Non | 210 | G/G | A/A | A/C | A/T | A/G | C/T | nd | nd | A/G | G/G | A/A | C/C | +/+ | +/+ | None | +/+ | +/+ | Present |
| Non | 212 | G/G | A/C | A/C | A/A | G/G | C/T | CAA/CAA | T/T | A/G | G/G | A/A | C/C | +/+ | +/+ | None | +/+ | +/+ | Present |
| Non | Kari | C/G | A/A | A/C | A/A | G/G | C/T | CAA/CAA | T/T | G/G | G/- | A/G | C/T | +/+ | +/+ | Het | +/+ | +/+ | Present |
| Ultramel | 250 | G/G | A/A | A/A | A/A | G/G | C/C | CAA/CAA | T/T | A/A | G/G | G/G | T/T | +/+ | +/+ | None | +/+ | *R305H / R305H* | Present |
| Ultramel | 259 | G/G | A/A | A/C | A/A | A/G | C/T | CAA/CAA | T/T | A/A | G/G | G/G | T/T | +/+ | +/+ | None | +/+ | *R305H / R305H* | Present |
| Ultramel | 550 | nd | nd | nd | nd | nd | nd | nd | nd | nd | nd | nd | nd | nd | nd | nd | nd | *R305H / R305H* | Present |
| Ultramel | 1025 | nd | nd | nd | nd | nd | nd | nd | nd | nd | nd | nd | nd | nd | nd | nd | nd | *R305H / R305H* | Present |
| Ultramel | 1263 | nd | nd | nd | nd | nd | nd | nd | nd | nd | nd | nd | nd | nd | nd | nd | nd | *R305H / R305H* | Present |
| Ultramel | 1376 | nd | nd | nd | nd | nd | nd | nd | nd | nd | nd | nd | nd | nd | nd | nd | nd | *R305H / R305H* | Present |
| Ultramel | 723 | nd | nd | nd | nd | nd | nd | nd | nd | nd | nd | nd | nd | nd | nd | nd | nd | *R305H / +* | Present |
| Ultramel | 872 | nd | nd | nd | nd | nd | nd | nd | nd | nd | nd | nd | nd | nd | nd | nd | nd | *R305H / +* | Present |
| Ultramel | 66 | G/G | A/A | A/C | A/T | A/G | T/C | nd | nd | A/G | G/- | A/A | C/C | +/+ | +/+ | None | +/+ | +/+ | Absent |
| Ultramel | 372 | nd | nd | nd | nd | nd | nd | nd | nd | nd | nd | nd | nd | nd | nd | nd | nd | +/+ | Absent |
| Ultramel | 1401 | nd | nd | nd | nd | nd | nd | nd | nd | nd | nd | nd | nd | nd | nd | nd | nd | +/+ | Absent |
